# The proline synthesis enzyme P5CS forms cytoophidia in *Drosophila*

**DOI:** 10.1101/2019.12.11.872267

**Authors:** Bo Zhang, Ömür Y. Tastan, Xian Zhou, Chen-Jun Guo, Xuyang Liu, Aaron Thind, Huan-Huan Hu, Suwen Zhao, Ji-Long Liu

## Abstract

Compartmentation of enzymes via filamentation has arisen as a mechanism for the regulation of metabolism. In 2010, three groups independently reported that CTP synthase (CTPS) can assemble into a filamentous structure termed the cytoophidium. In searching for CTPS-interacting proteins, here we perform a yeast two-hybrid screening of *Drosophila* proteins and identify a putative CTPS-interacting protein, Δ^1^-pyrroline-5-carboxylate synthase (P5CS). Using *Drosophila* follicle cell as the *in vivo* model, we confirm that P5CS forms cytoophidia, which are associated with CTPS cytoophidia. Overexpression of P5CS increases the length of CTPS cytoophidia. Conversely, filamentation of CTPS affects the morphology of P5CS cytoophidia. Finally, *in vitro* analyses confirm the filament-forming property of P5CS. Our work links CTPS with P5CS, two enzymes involved in the rate-limiting steps in pyrimidine and proline biosynthesis, respectively.

## Introduction

Cell metabolism manages energy mobilisation and utilization in each cell, by coordinating hundreds of thousands metabolic reactions occur simultaneously at any time. Spatial and temporal segregation of different reactions is critical for keeping the cell functionally normal. In eukaryotes, many membrane-bound organelles provide such a way to compartmentalise metabolic pathways. For example, mitochondrion is the place for oxidative reaction. Lysosomes house numeral digestive enzymes for the degradation of organelles and other molecules. Various post-translational protein modifications take place in the Golgi apparatus, while endoplasmic reticulum lies in the crossroad of protein synthesis.

However, the membrane-bound organelles do not account for the segregation of all metabolic reactions in the cell. The traditional view of cytosol as a homogeneous soup has recently been challenged. Many membraneless structures, such P bodies, purinosomes and U bodies have been identified in the cytoplasm (An et al., 2008; Liu and Gall, 2007; Sheth and Parker, 2006). In 2010, three studies independently reported that the metabolic enzyme CTP synthase (CTPS) can assemble into a filamentous structure termed the cytoophidium (which translates as cellular snake in Greek) in bacteria, yeast and fruit flies (Ingerson-Mahar et al., 2010; Liu, 2010; Noree et al., 2010). Subsequently, CTPS-containing cytoophidia can be found in human (Carcamo et al., 2011; Chen et al., 2011), fission yeast (Zhang et al., 2014) and plant (Daumann et al., 2018), suggesting filament-forming is an an evolutionarily-conserved property of CTPS (Liu, 2011, 2016). Additionally, genome-wide studies have shown that many more metabolic enzymes can form filamentous or dot structures in response to specific developmental stages or environmental stimuli (Noree et al., 2010; Shen et al., 2016).

CTPS catalyses the rate-limiting step of the *de novo* biosynthesis of CTP, an essential nucleotide for the synthesis of RNA, DNA and sialogycoproteins (Higgins et al., 2007). It also plays an important role in the synthesis of membrane phospholipids (Hatch and McClarty, 1996; McDonough et al., 1995; Ostrander et al., 1998). CTPS catalyses the ATP-dependent phosphorylation of UTP, followed by a glutaminase reaction that transfers the amide nitrogen to the C4 position of UTP to generate CTP (Lieberman, 1956; Long CW, 1967). CTPS activity is important to cell proliferation and has been demonstrated to be upregulated in cancers (Martin et al., 2014; Van Den Berg et al., 1995; van den BERG et al., 1993; Verschuur et al., 1998; Williams et al., 1978). Other studies have implicated abnormal CTPS activity with a number of human cancers, as well as with viral infection and parasitic diseases (De Clercq, 2001; Gharehbaghi et al., 2000; Kizaki et al., 1980; Verschuur et al., 2001; Verschuur et al., 2000; Weber et al., 1980).

Aiming to identify proteins interacting with CTPS, we carry out a genome-wide yeast two-hybrid screen and identify a putative CTPS-interacting protein, Δ^1^-pyrroline-5-carboxylate synthase (P5CS). P5CS, a bifunctional enzyme that encompasses simultaneously glutamate kinase and γ-glutamyl phosphate reductase activities, catalyses the reduction of glutamate to Δ^1^-pyrroline-5-carboxylate, which is subsequently converted to proline by P5C reductase (P5CR) (Hu et al., 1992; Smith et al., 1980; Vogel and Davis, 1952) (Chien-an et al., 1999; Merrill et al., 1989). P5CS controls the rate-limiting step in proline synthesis and negatively regulated by proline (Hong et al., 2000; Hu et al., 1992; Zhang et al., 1995). Defects in P5CS cause a connective tissue disorder characterized by lax skin and joint dislocations (Baumgartner et al., 2000; Baumgartner et al., 2005; Bicknell et al., 2008; Chien-an et al., 2008). In plant, P5CS is a stress-inducible gene and involved in salt and drought tolerance (Rai and Penna, 2013).

Here we identify P5CS as a novel filament-forming protein in *Drosophila* and reveal coordinated filamentation between P5CS and CTPS. Although these two enzymes have not been connected in previous biochemical studies, this study provides evidences supporting that P5CS and CTPS are coordinated spatially.

## Results

### P5CS forms cytoophidia in *Drosophila* cells

Using the full-length *Drosophila* CTPS as a prey, we carried out a yeast two-hybrid analysis by screening a genome-wide library of *Drosophila* peptides for potential interacting partners for CTPS. To this end, we identified the positive protein, the product of gene CG7470 could interact with CTPS. The gene *CG7470* codes for a 776-aa protein. Bioinformatics analysis revealed that CG7470 is a *Drosophila* orthologue of delta1-pyrroline-5-carboxylate synthetase (P5CS). These results caught us a surprise since the connection between CTPS and P5CS has not been revealed in previous biochemical studies.

It is known that CTPS forms filamentous cytoophidia in various tissues in *Drosophila*, especially in the female reproductive system (Liu, 2010). To test if P5CS forms similar filamentous structures, we generated transgenic flies carrying the Venus-P5CS genetic information and dissected adult flies expressing Venus-P5CS. We observed that Venus-P5CS forms filamentous cytoophidia in follicle cells (Figure 1A and 1B). More specifically, Venus-P5CS forms linear cytoophidia in some cells, while the cytoophidia curl up to form ring-shaped structures in other cells. Very frequently, we observed linear cytoophidia with a small ring-structure at one end.

**Figure 1.**
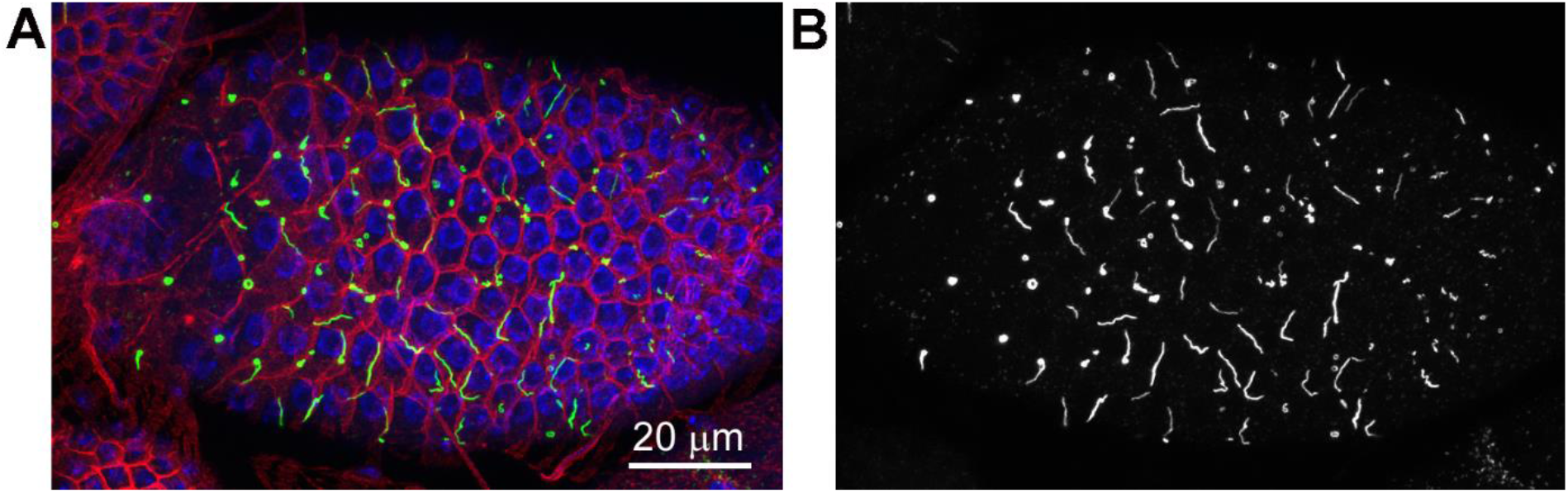
Venus-P5CS forms filamentous structures in *Drosophila* follicle cells. Generation of transgenic fly with Venus-P5CS. (A) Confocal images of Venus-P5CS filaments (green) in follicle cells of a stage 9 egg chamber. DNA is labelled with Hoechst 33342 (blue). Cell membrane is labelled with Hu-li tai shao (red). Scale bar, 20μm. (B) Representative linear or curly P5CS filaments and ring-shaped structures at one end of linear filaments are shown.

To study the expression profile of P5CS in *Drosophila*, we quantified its mRNA level in different developmental stages and different tissues in *Drosophila* by qPCR. During embryogenesis, P5CS mRNA reached its highest levels in the 4-8 h embryos and was about 9-fold to 0-4 h embryos. In embryos at 8-20 h and 20-24 h, P5CS stayed nearly 6-fold (Figure S1A). In larval stages, P5CS showed the strongest expression in first instar larvae and decreased to 0.25- and 0.4-fold in second and third instar larvae, respectively (Figure S1B). P5CS expression level increased during pupal development and reached its peak in the late pupal stage (Figure S1C). In adult flies, P5CS showed low expression level in ovary, while abundant expression was detected in heads and gut, with the highest level in males (Figure S1D). Then, we extended our studies to other tissues of *Drosophila melanogaster*. We found P5CS filamentation in all the tissues we examined, including wind discs, ventral nerve cords, larval midguts, salivary glands, tracheae, adult hindguts, accessory glands and ejaculatory ducts (Figure 2A-2H).

**Figure 2.**
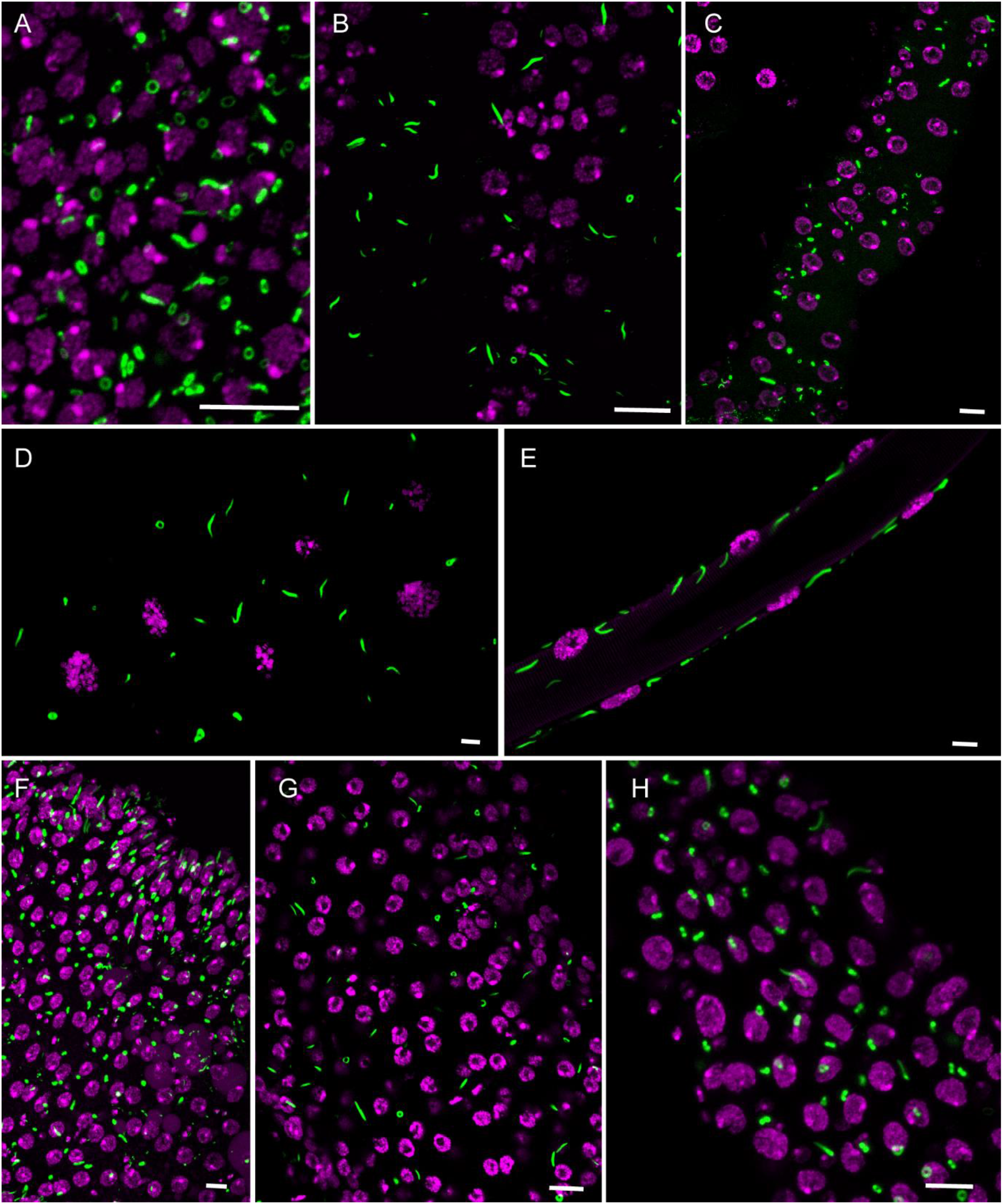
P5CS forms cytoophidia in multiple tissues. Tissues were derived from offsprings of *UAS-Venus-P5CS* crossed with *da-GAL4* lines. DNA is stained with Hoechst 33342 (magenta). (A) Wing disc. (B) Larval ventral nerve cord. (C) Larval midgut. (D) Larval salivary gland. (E) Adult trachea. (F) Adult hindgut. (G) Adult accessory gland. (A) Adult ejaculatory duct. Scale bars, 10 μm.

To address the concern that Venus tag might promote filamentation of P5CS artificially, we generated transgenic flies expressing P5CS tagged with HA, a tag much smaller than Venus. Using antibodies against HA, we were able to detect HA-P5CS filaments in *Drosophila* follicle cells (Figure S2).

### Association of P5CS and CTPS cytoophidia

Glutamate, the product of CTPS catalytic reaction, serves as the substrate of P5CS. Since both CTPS and P5CS form filamentous structures, we sought to understand the association between P5CS and CTPS cytoophidia. We speculated that there are three possibilities regarding the relationship of these two kinds of cytoophidia.

First, both CTPS and P5CS may be components of the same structure, having a relationship similar to that between alpha-tubulin and beta-tubulin. We refer to this type of relationship as ‘dependent filamentation’. In this case, we would expect P5CS and CTPS to show identical morphology and distribution under light microscopy. Furthermore, removal of one component would disrupt the filamentation of the other protein. Changing the levels of one protein would impact the distribution of the other.

The second possibility is the case that we refer to as ‘independent filamentation’. In this occasion, the P5CS cytoophidium is independent from the CTPS cytoophidium and vice versa. In terms of localisation, P5CS and CTPS should not colocalise with each other. Disruption of one type of filament should have no effect on the other type. Moreover, overexpression of one type of filament should not affect the other.

There is a third possibility, to which we refer as ‘interdependent filamentation’ or ‘coordinated filamentation’. In this scenario, the distribution of the two types of cytoophidia would be similar, but not identical, to each other. Unlike ‘dependent filamentation’, ‘coordinated filamentation’ should not abolish the filamentation of one kind when the other is disrupted. Changing one kind of cytoophidia in the case of ‘coordinated filamentation’ would affect the other kind, in contrast to the situation of ‘independent filamentation’.

To determine the relationship of P5CS and CTPS, we stained follicle cells from Venus-P5CS flies with an antibody against CTPS. Using confocal microscopy, we observed that Venus-P5CS and CTPS showing similar, but not identical distributions (Figure 3A). More specifically, P5CS cytoophidia were curly in shape and often formed a small ring at one end, whereas CTPS cytoophidia were straight and without a curly end (Figure 3B). While for most of the length of P5CS cytoophidia there was a colocalisation with CTPS cytoophidia, we did not detect a CTPS signal on the curly end of P5CS cytoophidia. Therefore, merged images of CTPS (in red) and P5CS (in green) show yellowish stems in connection with the curly green end (Figure 3B). Quantification of the signal intensities of CTPS and P5CS along the long axes of the cytoophidia clearly demonstrated the differences in the localisation of these two filaments (Figure 3C). P5CS and CTPS showed correlated (neither identical nor random) distributions, suggesting that filamentation of these two enzymes is interdependent and coordinated.

**Figure 3.**
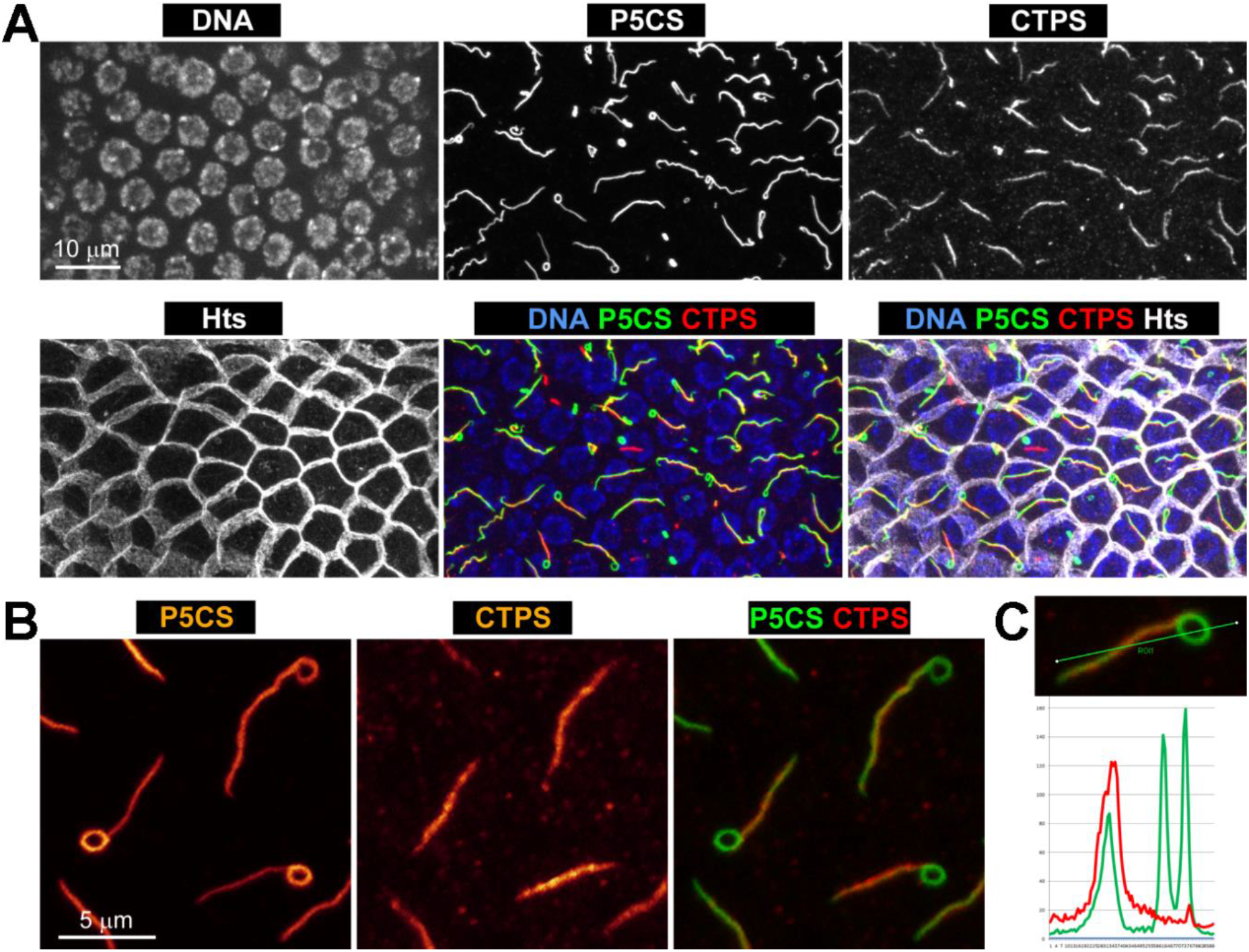
Association of P5CS and CTPS cytoophidia in *Drosophila* follicle cells. (A) Immunostaining results of P5CS, CTPS and merged picture are presented. The yellowish indicated the overlap bulk of CTPS and P5CS filaments. Scale bar, 10 μm. (B) Zoom-in views of some parts of CTPS, P5CS filaments and merged results from (A) are shown. The green ring-shaped structure suggests that it contains only P5CS filament. Scale bar, 5μm. (C) Fluorescence intensity of overlapping part of two types of filaments and the ring-shaped structure of P5CS filament is measured. Green, P5CS signal. Red, CTPS signal.

To better understand the distribution of P5CS and CTPS between cells, ovaries from Venus-P5CS flies were stained by antibodies against CTPS and Hu-li tai shao, a membrane protein (Figure 4A). The signal of Hu-li tai shao outlined the boundary of follicle cells. We observed that P5CS forms long filaments spanning multiple cells. One or both ends of P5CS filaments can be anchored on the cell cortex. On the contrary, CTPS cytoophidia were constrained inside individual follicle cells (Figure 4B and 4C). These results further demonstrated that P5CS and CTPS cytoophidia are not parts of the same structure even though they localise adjacently to each other. In addition, we observed that two P5CS filaments intertwine with one CTPS filament (Figure S3).

**Figure 4.**
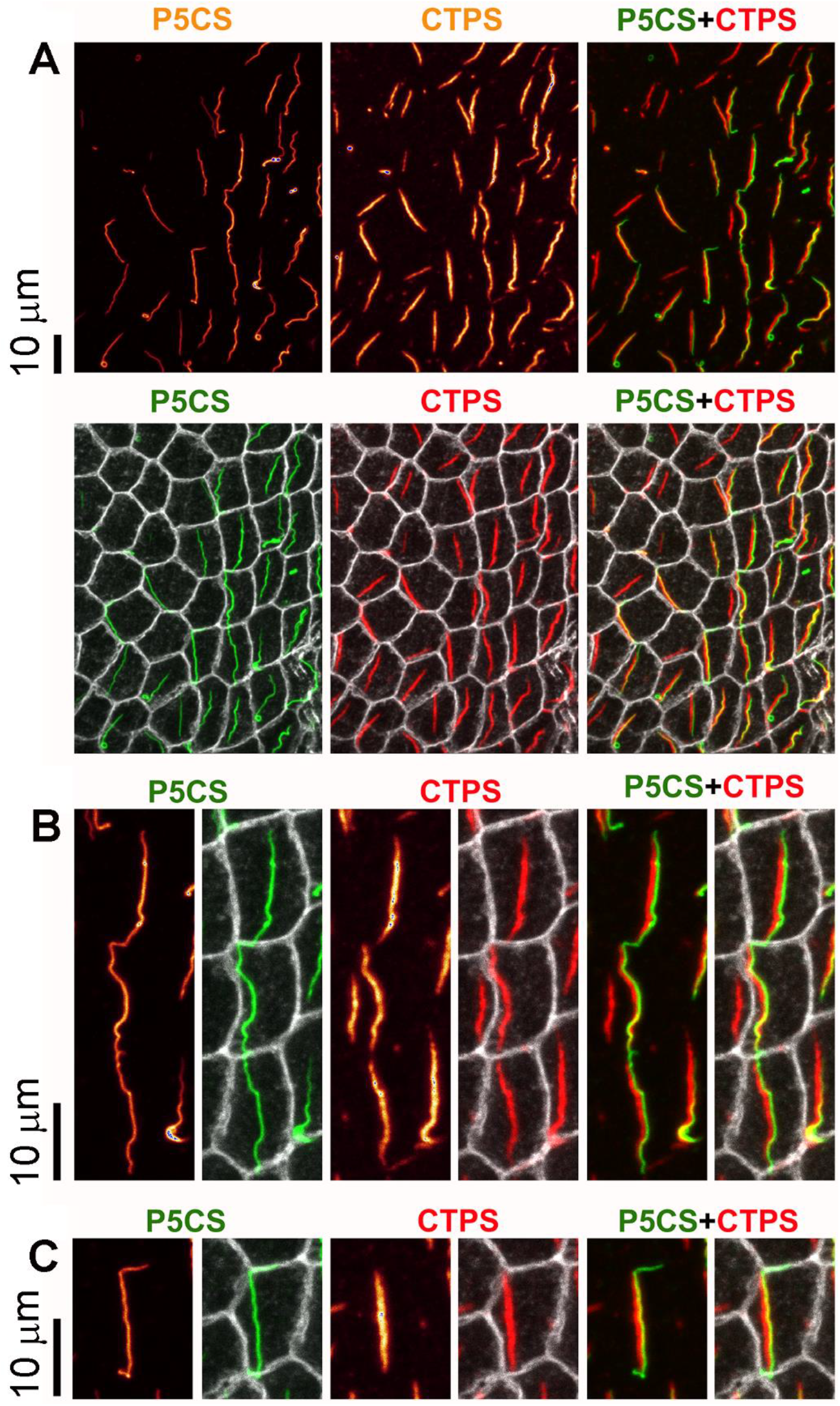
P5CS cytoophidia are anchored on cell cortex. (A) Confocal images of P5CS, CTPS filaments and merged results (cell membrane, white). (B) The relationship between P5CS, CTPS filaments and cell membrane is shown. (C) Zoom-in view of (A). Scale bar, 10 μm.

### Coordinated filamentation of P5CS and CTPS

Next, we sought to investigate how CTPS influences P5CS filamentation. As we described above (Figure 3B), Venus-P5CS filament has a curly end in which CTPS is undetectable. We hypothesise that CTPS affects the curvature of P5CS filaments. We predict that if the CTPS filament becomes more abundant, P5CS would become straighter, whereas disrupting CTPS filaments will promote the curvature of P5CS filaments.

In our previous study, we found that overexpressing a truncated version of CTPS (i.e. having only its synthetase domain) has a dominant-negative effect on CTPS filamentation (Azzam and Liu, 2013). This means that expressing CTPS synthetase domain ectopically prevents filamentation of endogenous CTPS. Having this knowledge, we expressed CTPS synthetase domain ectopically to disrupt CTPS filamentation in a Venus-P5CS background. As expected, we no longer detected clear CTPS filaments in follicle cells as compared with the control group (Figure 5A and 5C). When CTPS filaments were disrupted, we observed that Venus-P5CS formed ring-shaped structures instead of straight filaments (Figure 5B and 5D). The abundance of ring-shaped P5CS structures increased significantly when CTPS filamentation was disrupted. Quantification results showed that the curvature of P5CS filaments increased while the overall length decreased significantly (Figure 5G and 5H). These data suggest that CTPS filaments stabilise or straighten P5CS filaments.

**Figure 5.**
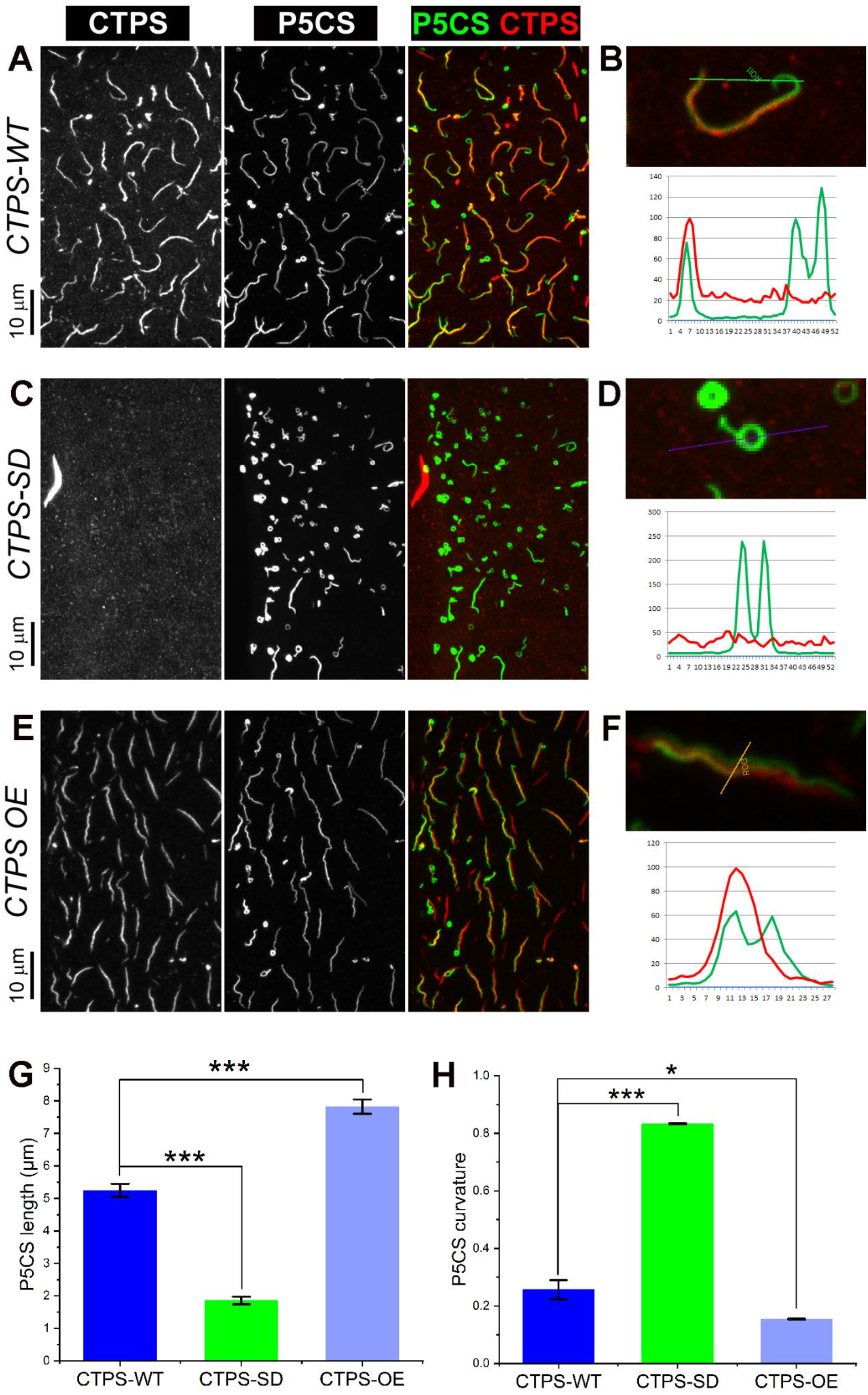
CTPS filamentation affects the morphology of P5CS cytoophidia. (A), (C), (E) Conformational results of CTPS, P5CS filaments and merged images under CTPS cytoophidia wild-type (A), disruption (C) and overexpression (E) levels in follicle cells. Scale bar, 10μm. (B), (D), (F) Fluorescence intensity of P5CS and CTPS filaments is measured under different CTPS levels. (G), (H) Quantification analysis of length (G) and curvature (H) of P5CS filaments under different CTPS levels. *P<0.05, ***P<0.001, Error bars show the SEM.

If the hypothesis that CTPS filaments help the straightening of P5CS filaments is correct, we would expect that increasing CTPS filaments have the opposite effect (i.e. making P5CS filaments longer and less curly). Our previous study shows that overexpressing CTPS induces the formation of large and long filaments. Indeed, we found that P5CS filaments in cells overexpressing CTPS became longer and less curly than those in control cells (Figure 5E-5H). These data support the idea that the filamentation of P5CS and CTPS is coordinated.

### P5CS effects on CTPS filamentation

To further investigate the role of P5CS on CTPS cytoophidium formation, we analysed the effect of P5CS knockdown using RNAi in stage 9 follicle cell clonal patches. RT qPCR analyses showed that all three P5CS RNAi lines (v10176, v38955 and v38953) significantly reduced the P5CS expression in comparison to the mCherry control, whilst the three P5CS lines had no effect on CTPS expression (Figure S4A and B). Using an inducible Tub<GAL80>GAL4 driver, the mosaic expression of the three P5CS RNAi lines was monitored by the expression of GFP in cell nuclei (Figure S4C, D, F, G, I and J). Expression of these three P5CS RNAi lines had no significant effect on the length or width of CTPS cytoophidia, indicating that P5CS is not necessary for the CTPS filamentation (Figure S4E, H and K). These data support the idea that P5CS and CTPS cytoophidia are not in the same structure.

Then we wondered whether the length or width of CTPS cytoophidia would change upon overexpression of P5CS. Both UAS-mCherry control and UAS-Venus-P5CS flies were crossed with Actin5c-GAL4 driver. CTPS cytoophidia appeared visibly larger in follicle cells overexpressing Venus-P5CS (Figure 6A and 6B). Quantification showed that the area of CTPS cytoophidia in P5CS overexpression background was 1.5 fold greater compared to the mCherry control (Figure 6C). These data argue against the idea that CTPS filamentation is independent of P5CS.

**Figure 6.**
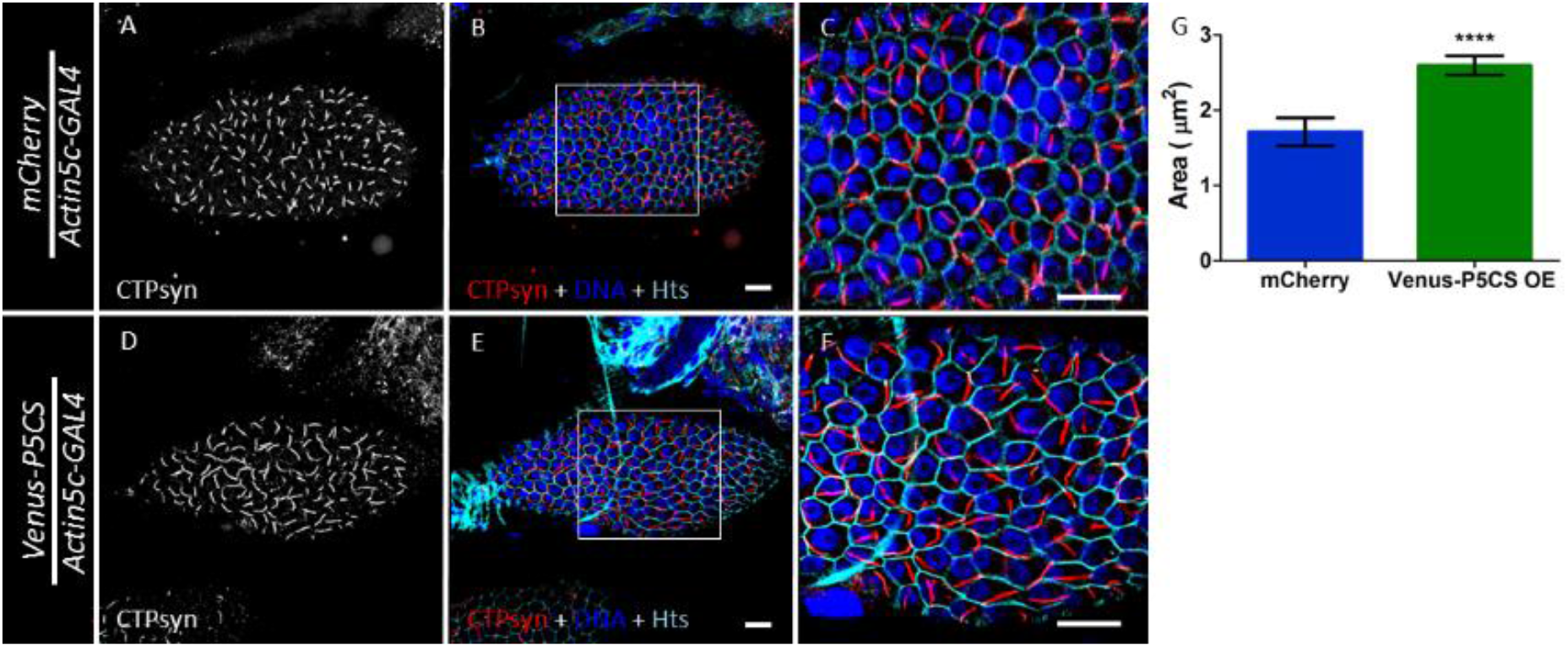
Overexpressing P5CS increases the length of CTPS cytoophidia. Representative images of CTPS filaments in mCherry control (A and B) and P5CS overexpression lines (D and E). Both UAS-mCherry control and UAS-Venus-P5CS flies were crossed with Actin5c-GAL4 driver. (C), (F) Zoom-in views of (B) and (E), respectively. Scale bars, 10 μm. (G) CTPS cytoophidium areas of mCherry control and P5CS overexpression line are quantified. ****P<0.0001, Error bars show SEM.

### Mutations in P5CS dimerisation interface impede filament formation

The human *P5CS* gene, known as *ALDH18A1*, plays an essential role in the interconversion of glutamate, ornithine and proline (Hu et al., 2008; Perez-Arellano et al., 2010). The deficiency of P5CS activity in humans is associated with a rare, inherited metabolic disease (Perez-Arellano et al., 2010). Human P5CS possesses two enzymatic domains, γ-glutamyl kinase (1-361aa) and γ-glutamyl phosphate reductase (362-795aa) (Hu et al., 2008). We aligned the protein sequences of *Drosophila* and human P5CS and found that they are more than 60% identical. Here we built a 3D model of *Drosophila* P5CS using the human P5CS structure as a reference. The homology model of *Drosophila* P5CS shared the same dimer interface and conserved residues in the dimerisation interface, including L450, N708, F713, H755 and F760 (Figure 7A and 7B). We constructed different mutants in the dimerisation interface and transfected them into S2 cells. Using Venus-P5CS as a control, we found that two of the mutants, N708 and F713, could impede P5CS filament formation (Figure 7C). The signal of Venus-P5CS^N708A^ exhibited a diffused localisation pattern and Venus-P5CS^F713A^ formed dotted structures in cells.

**Figure 7.**
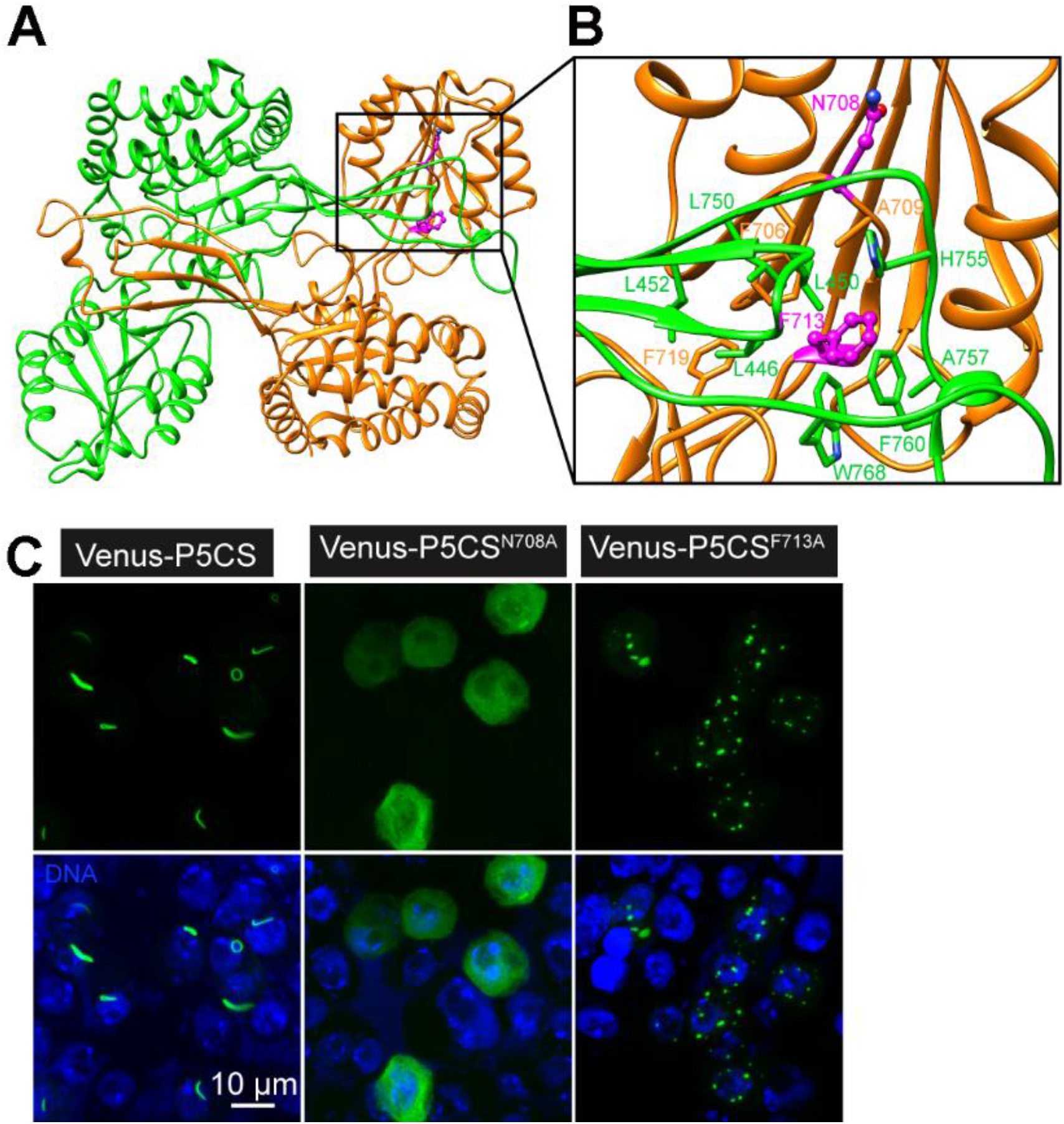
P5CS dimerisation interface mutants disrupt filament formation. (A) Homology model of P5CS dimer with human P5CS structure as reference. (B) P5CS dimer interface is presented. Key residues in dimerisation interface are labelled and mutation sites that may have a strong effect on dimerisation of P5CS are highlighted in blue (N708 and F713). (C) Venus-P5CS, Venus-P5CS^N708A^ and Venus-P5CS^F713A^ were cloned into pAc 5.1 vector and transfected into S2 cells. Representative confocal images of cells are presented. P5CS signal is shown in green. DNA is labelled with Hoechst 33342 (blue). Scale bar, 10 μm.

### P5CS exhibits filament-forming capability *in vitro*

To determine if P5CS can form filaments *in vitro*, we expressed *Drosophila* P5CS in *E. coli* cells and purified the protein. We used electron microscopy to analyse the filament-forming capability of P5CS *in vitro* under various conditions (Figure 8).

**Figure 8.**
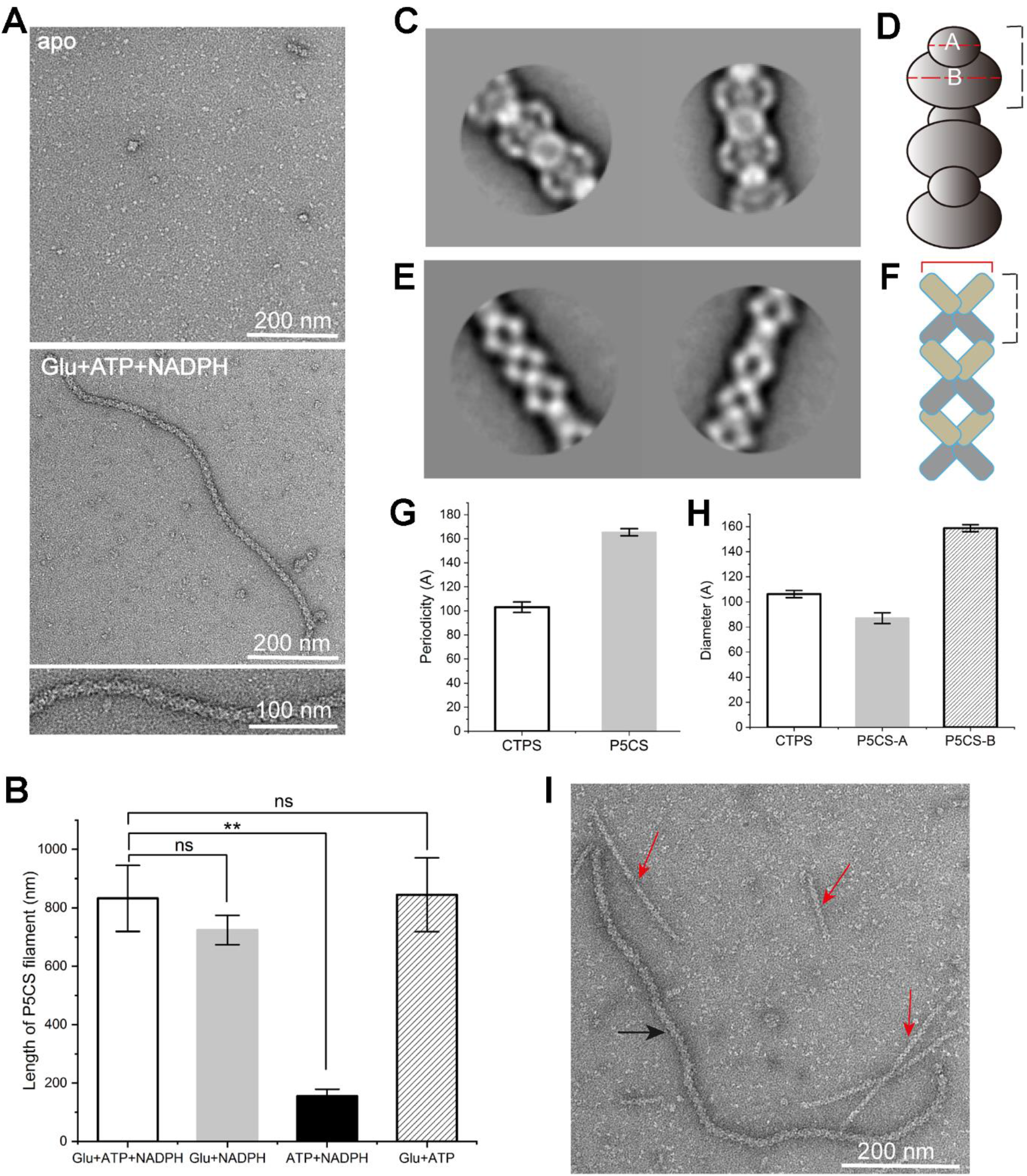
P5CS forms filamentous structures *in vitro*. (A) Negative stain electron microscopy images of P5CS conformation in apo and substrate-bound states (Glutamate, ATP and NADPH). A magnified filament image is shown in lower panel. (B) Quantification analysis of filaments length in different conditions by removing glutamate, ATP and NADPH successively from the complete reaction. **P<0.01. Error bars show the SEM. (C-F) 2D classification of two kinds of filaments. (G and H) Quantification and comparison between the two types of filaments on their periodicity and diameter. Error bars show the SEM. (I) A representative image of filaments in the condition with substrates of CTPS catalytic reaction (Glutamine, ATP, UTP and GTP) and substrates of P5CS catalytic reaction (Glutamate, ATP and NADPH). Two kinds of filaments are indicated by red and black arrows.

Provided with ATP and NADPH, the metabolic enzyme P5CS catalyzes the conversion of glutamate into Δ^1^-pyrroline-5-carboxylate (Fujita et al., 1998; Ginzberg et al., 1998; Hu et al., 1992). Purified P5CS in apo state could hardly form filaments. By contrast, when its substrates (ATP, NADPH and Glutamate) were provided, P5CS could form long filaments after a 10-minute incubation at 25°C (Figure 8A). Removing ATP or NADPH from the solution did not prevent filamentation of P5CS, suggesting that the filament formation is not depended on the reaction *per se*. However, removing glutamate from the solution almost abolished P5CS filament formation (Figure 8B). These results demonstrate that *Drosophila* P5CS has filament-forming capability *in vitro*.

When both CTPS and P5CS were incubated in the same test tube, we observed that P5CS could form filaments independent of the CTPS filaments (Figure 8I and S5). The 2D classification of P5CS filaments is distinct from CTPS filaments (Figure 8C and 8E). The basic unit of the P5CS filaments is bow-tie shaped (Figure 8D), while the basic unit of the CTPS filaments is X-shaped (Figure 8F). The diameter of the P5CS filament is 159Å with a periodicity of 165Å, while the diameter of the CTPS filament is 106Å with a periodicity of 103Å (Figure 8G and 8H).

## Discussion

Using the *Drosophila* follicle cell as a model system, we demonstrate that CTPS and P5CS form cytoophidia interdependently. Both CTPS and P5CS cytoophidia show very similar patterns. However, we provide evidence to the fact that they are not the same structure. First, we observe that P5CS filaments entangle with CTPS filaments. Image analysis shows that these strings are very close but not identical. Second, the ends of P5CS filaments frequently associate with cell cortex, whereas CTPS does not show clear association with cell cortex. Third, in some cases we can see P5CS filaments continuously crossing from one cell to the neighbour cell or even further, whereas CTPS filaments are constrained inside individual cells. Fourth, although P5CS and CTPS intertwine, P5CS filaments tend to be curlier than CTPS filaments.

There are several hypothetical advantages for the storage of enzymes such as CTPS and P5CS in filamentous form. Polymerisation may play a role in controlling enzyme activity in response to demand through stabilisation of the enzyme in active or inactive states. It is unclear whether filament-forming enzymes are subject to enzymatic downregulation or upregulation when assembled into cytoplasmic filaments. For example, acetyl CoA carboxylase (ACC) activity is upregulated when polymerised (Kim et al., 2010), whilst glutamine synthase activity is downregulated upon polymerisation. The storage of metabolic enzymes in filaments may provide rapid changes to enzyme activity in response to changes in cellular metabolic needs. Furthermore, the restriction of the metabolic enzymes through filament formation to certain subcellular areas may function in such a way that a concentration gradient of substrates and products within the cytosol is created and maintained.

Why filamentation of P5CS needs to be coordinated with that of CTPS? A potential reason for coordinated filamentation between these two enzymes could be that the close association of the enzymes might enable metabolic channelling. The product of one enzyme, not being released into solution, can pass directly onto another enzyme. For example, glutamate, a product of CTPS, serves as a substrate for P5CS and may also regulate filamentation of P5CS.

There are a number of advantages of metabolic channelling over the free diffusion of reaction products. Firstly, metabolic channelling makes a metabolic pathway more efficient that diffusion as the transit time from one active site to the next is reduced. Secondly, it protects the intermediate product from decomposition by the aqueous external environment. Thirdly, channelling may segregate substrates and products from competing enzymatic reactions and circumvent unfavourable equilibria.

Metabolic channelling, however, does not explain why P5CS and CTPS do not always colocalise with each other. Since filament formation can regulate enzymatic activity, areas where P5CS and CTPS do not overlap may represent a depot for inactive enzyme where metabolic channelling is not required. Independent filaments could be induced as a result of storage of excess P5CS.

In summary, this study identifies the filament-forming property of P5CS both *in vivo* and *in vitro*. Coordinated filamentation of P5CS and CTPS provides a mechanism for intracellular compartmentation without membrane, extending our understanding of the complexity of cellular organisation. The close relationship between CTPS and P5CS demonstrated in this work has escaped detection from previous biochemical studies, highlighting the importance of investigating metabolic compartments *in situ*.

## Materials and Methods

### Yeast two hybrid screen

A yeast two-hybrid screen was carried out by Hybrigenics Services (Cambridge, MA, USA). The full-length *Drosophila melanogaster* CG6854 protein was used as bait. The screen was performed on *Drosophila* Whole Embryo cDNA library using two different fusions of N-LexA-CG6854-C and N-Gal4-CG6854-C. The screen identified P5CS as an interacting protein with CTPS.

### *Drosophila melanogaster* stocks and genetics

All stocks were raised at 21°C on standard cornmeal. The stocks used were: *hsFLP;UAS-GFPnls;UAS-dcr2; tub<Gal80>Gal4/SM5, Cy-TM6,Tb* (inducible Tub-GAL4 driver stock), *w;Act5cGAL4/Cyo twi 2xEGFP* (Actin5c-GAL4 stock), *UASp-Venus-CG7470/cyo* (UAS-Venus-P5CS transgene stock), *UAS-mcherry.VALIUM10* (UAS-mCherry stock), UAS-CTP synthase^JF02214^ (CTPsyn RNAi stock). Three P5CS RNAi stocks were used, stock number: v101476, v38953 and v38955.

### Transgenic flies

The Venus-P5CS transgene was generated using the P5CS cDNA clones received from DGRC (clone number: GH12632). The cDNA was cloned into an entry vector by TOPO cloning and then Gateway cloning was employed to clone into a *Drosophila* Gateway™ Vector Collection destination vector, pPVW. To overexpress the Venus tagged P5CS transgene ubiquitously in flies, they were crossed to Actin5c-GAL4 flies and recombinants were generated.

### Total RNA extraction and reverse transcription

For quantification of the RNAi knockdown, UAS-mCherry and RNAi lines were crossed with an *Actin5c-GAL4* driver, and at least 3 samples of 20 first instar larvae progeny from each group were collected and washed with PBS. Samples were homogenized using the Qiagen QIAshredder (Cat. no. 79654) and RNA was extracted using the Qiagen RNeasy Plus Mini Kit (Cat. No. 74134) as per the manufacturer’s instructions. RNA samples were kept at −80°C. Using the Qiagen QuantiTect Reverse Transciption kit (Cat. no. 205311), reverse transcription was carried out on 500ng RNA, following the manufacturer’s instructions including the genomic DNA removal step. The resulting cDNA was diluted 1:10 using nuclease-free water and kept at −20°C.

### Quantitative PCR (qPCR)

1μl of the cDNA from the reverse transcription stage was mixed with 5μl 2x SYBR^®^ Green JumpStart™ Taq ReadyMix™ (Sigma Aldrich), 0.8μM of primer (Actin5c, CTPsyn, P5CS) and diluted with H2O for each 10μl qPCR reaction. The reactions were carried out using the 7500 Fast RealTime PCR System (Applied Biosystems) on the normal setting: initial denaturation at 95°C for 2 minutes, followed by 40 cycles of 15sec denaturation at 95°C, 30sec primer annealing at 55°C, 30 seconds elongation at 72°C. Expression values were normalised using reference gene *Actin5c*.

### Immunohistochemistry

*Drosophila* ovaries, were dissected in Grace’s medium (Invitrogen Cat. no. 11605045) and fixed with 4% paraformaldehyde for 10 min and then washed 3 times with PBT (PBS+0.4% Triton X-100) for 2 minutes each time. Samples were incubated with primary antibody overnight at room temperature. They were then washed and incubated with DNA dye Hoescht 33342 (1:10000) and secondary antibodies overnight at 4°C. Primary antibodies used in this study were rabbit anti-CTPsyn (1:1000; y-88, sc-134457, Santa Cruz BioTech Ltd, Santa Cruz, CA, USA), mouse anti-Hu-li tao shao (Hts) (1:20; 7H9 1B1, Developmental Studies Hybridoma Bank, Iowa City, IA, USA). Secondary antibodies used in this study were donkey anti-mouse and anti-rabbit antibodies that were labeled with Alexa Fluor^®^ 488 and Cy5, respectively (1:500, Jackson ImmunoResearch Laboratories, Inc., West Grove, PA, USA).

### Laser-scanning confocal microscopy

3D-stacks of stage 9 egg chamber images were acquired under a 63 × oil objective on laser-scanning confocal microscopes (Leica SP5 or SP8 Confocal Microscope). For quantification of cytoophidia length and width in follicle cells, length and width of cytoophidia from 30 stage 9 egg chambers from each fly line were measured using the ‘analyse particles’ tool in ImageJ (v1.43 U). As cytoophidia in the ovaries expressing the Venus-P5CS transgene were not straight, the length of cytoophidia couldn’t be measured accurately, so the area was measured instead.

### Protein expression and purification

*Drosophila* CTPS and P5CS were cloned into pET28a vector with a C-terminal 6XHis and N-terminal 6XHis-SUMO tag separately. All vectors were transformed into Transetta (DE3) E. coli cells. Followed by induction with 1 mM isoprol β-d-1-thiogalactopyranoside (IPTG) when OD600 reached 0.8, proteins were expressed at 16°C for 16~18 hours. The cells were harvested by centrifugation and resuspended in pre-cold lysis buffer (50 mM Tris-HCl, pH 8.0, 500 mM NaCl, 10% glycerol, 10 mM imidazole, 5 mM β-Mercaptoethanol, 1mM PMSF, 5 mM Benzamidine Hydrochloride). Subsequently, the cells were lysed with ultrasonic cell disruptor and centrifuged. The supernatant was collected and incubated with equilibrated Ni-NTA Agarose (30250; Qiagen) beads at 4°C for 1 hour. Proteins were washed with ice cold lysis buffer and then eluted with elution buffer (50 mM Tris-HCl, pH 8.0, 500 mM NaCl, 10% glucerol, 250 mM imidazole, 5 mM β-Mercaptoethanol). CTPS-6X His protein was concentrated to 2mL. 6XHis-SUMO-P5CS was cleaved by ULP1 at 4°C overnight and then concentrated to 2mL. Concentrated proteins were loaded to size-exclusion chromatography with Hiload 16/600 Superdex 200 pg (28989335; GE Healthcare). Fractions were collected and analysed by SDS/PAGE. Fractions containing *Drosophila* CTPS or P5CS were concentrated and stored in storage buffer (20 mM Tris-HCl, pH 8.0, 150 mM NaCl, 10% glycerol).

### Electron microscopy

Samples for negative stain electron microscopy were prepared by applying P5CS to carbon-coated grid and staining with 2% uranyl acetate. 300 μM P5CS in 20 mM Hepes (pH 8.0) and 10 mM MgCl2, supplemented with 30mM Glutamate, 2 mM ATP, 0.5 mM NADPH, or removing one from the substrates, or without any substrates as a control, was incubated for 10 minutes at 25°C before being coated onto grids. For CTPS and P5CS mixed reaction, samples were prepared with 300 μM P5CS and 300 μM CTPS in 20 mM Hepes (pH 8.0), supplemented with 10 mM MgCl2, 10mM Glutamine, 2 mM ATP, 2mM UTP, 0.2 mM GTP, 30mM Glutamate and 0.5 mM NADPH, and samples were incubated for 10 minutes at 25°C. Negative stain EM was performed on a Tecnai G2 Spirit (FEI co.) operating at 120 kV, and images were acquired at 52,000X magnification on a US4000 4K X 4K CCD Camera (Gatan Inc.).

### Statistical analysis

Raw data was entered into Prism (v7.00, GraphPad, CA) and used to produce graphs. The results are shown as mean values ± standard deviation. Before any statistical analysis, data was confirmed to be normal. To test the significance of cytoophidia size compared to wild-type controls, a two-tailed Student t test was performed unless otherwise specified. For data involving more than two groups, a one-way ANOVA test was performed, followed by Dunnett post-hoc test to check for significant differences between data groups. Significant differences were attributed for p<0.05.

## Acknowledgments

We are grateful to Andrew Bassett and Mayte Siswick for technical support and to Christos Andreadis and Chia Chun Chang for reading manuscript. We thank Ying Han for assistance with mass spectrometry equipment, Xiaoming Li for training on use of the confocal microscope and software guidance, Tiezhu Shi for discussion on data analysis. We also thank Bio-Electron Microscopy Facility of ShanghaiTech University for providing assistance.

## Supporting information (Figures S1-S5)

**Figure S1.**
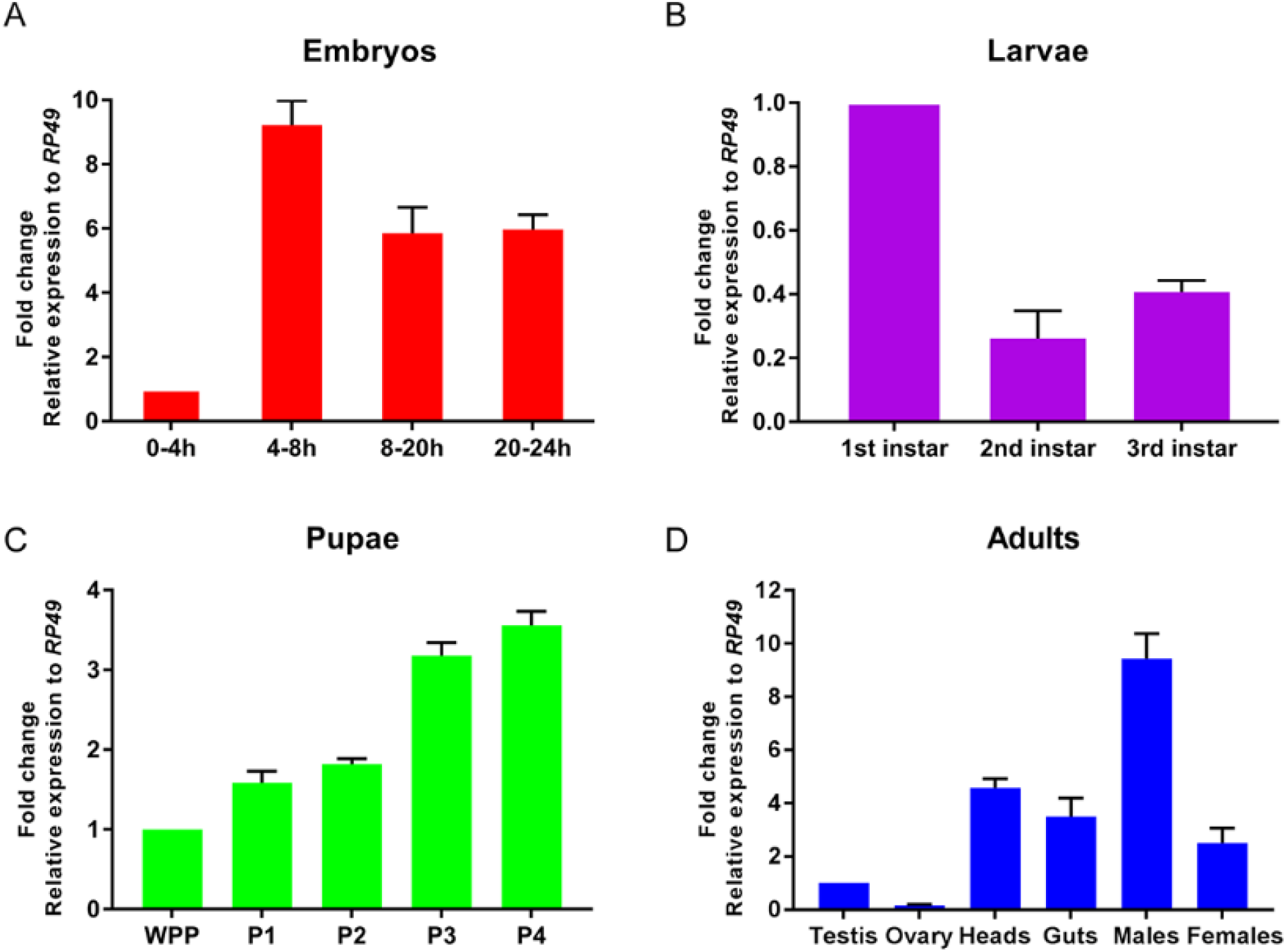
Expression profile of *Drosophila* P5CS as revealed by qPCR. (A-D) P5CS expression levels in embryos, larvae, pupae and adults, respectively. The fold change of 0-4h, 1st instar larvae, WPP and testis was normalized to 1 during different developmental stages, respectively. All flies are from *w^1118^*. WPP, white pre-pupae. P1-P4, pupal stages 1-4.

**Figure S2.**
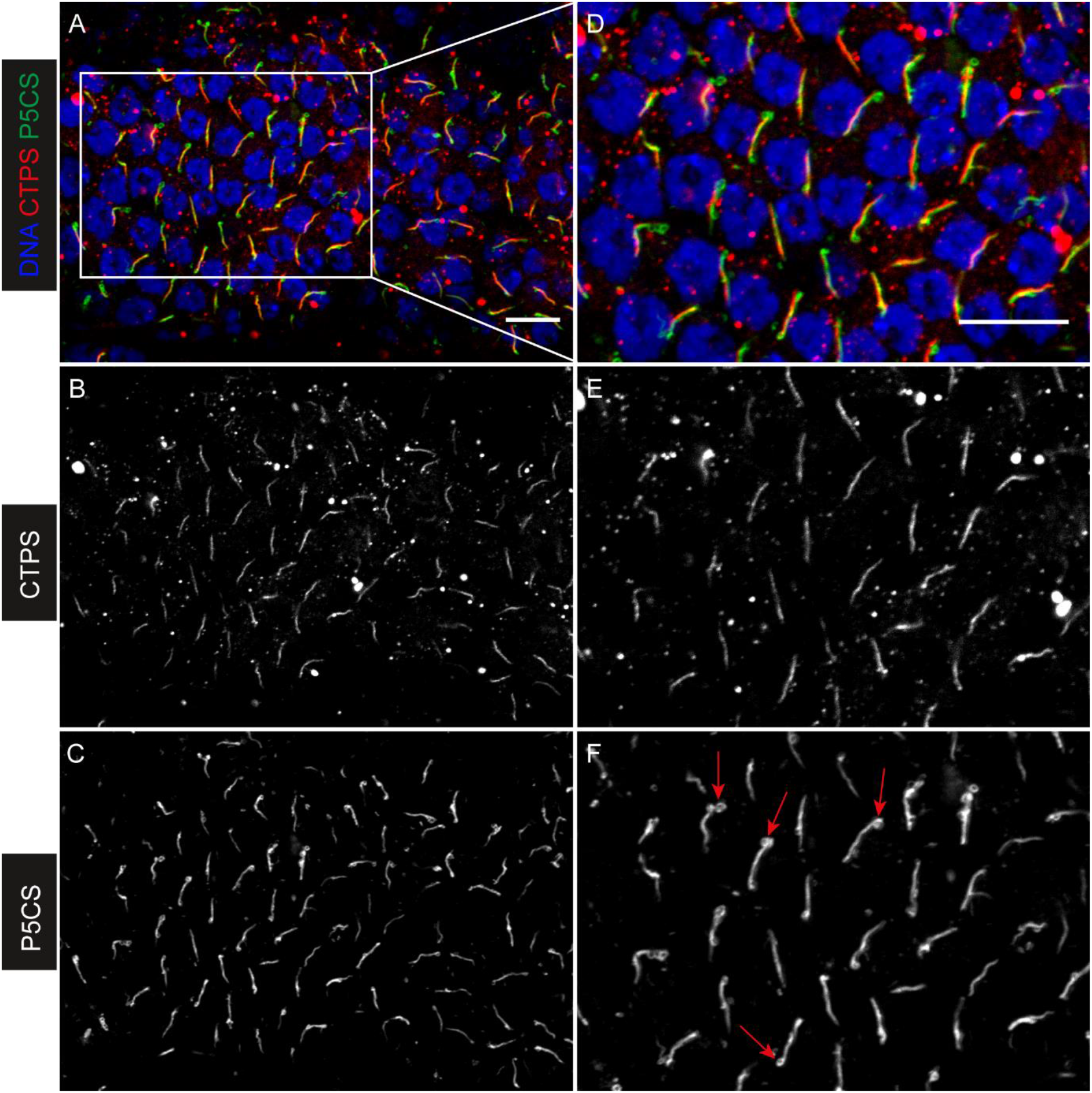
HA-P5CS forms filaments in *Drosophila* follicle cells. (A) Both CTPS (red) and HA-P5CS (green) form filaments in follicle cells. DNA is labelled with Hoechst 33342 (blue). (B) CTPS only. (C) HA-P5CS only. (D-F) Zoom-in images of (A-C), respectively. Ring structures of HA-P5CS are indicated by red arrows in (F). Scale bars, 10μm.

**Figure S3.**
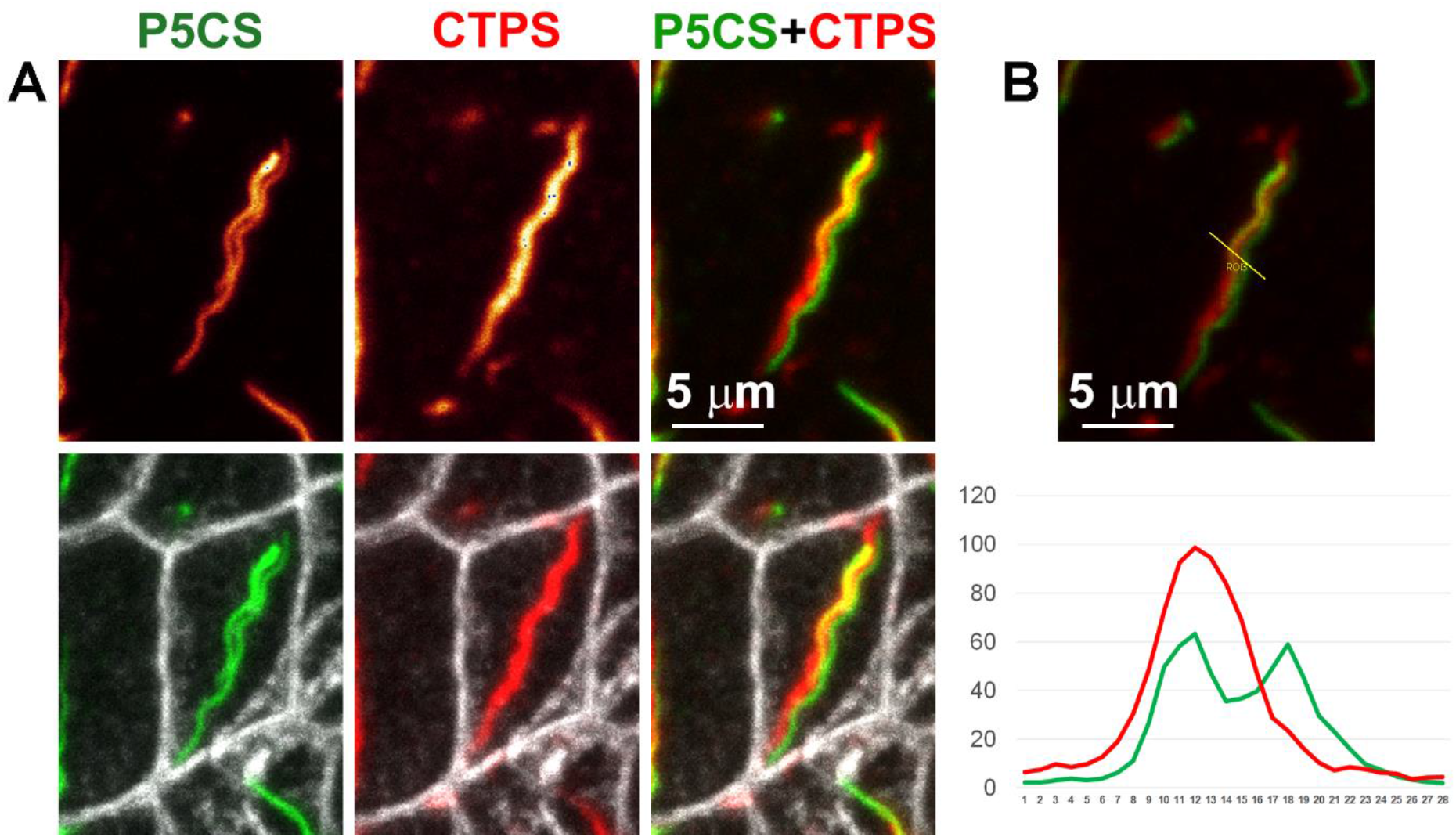
Two P5CS filaments intertwine with one CTPS filament. (A) Two P5CS filaments (green) intertwine with one CTPS filament (red) in *Drosophila* follicle cells. The outline of follicle cells was labelled with a membrane protein Hu-li tai shao (white in low row images). Note this is a zoom-in image shown in Fig 5E. (B) The intensity analysis of the cross section indicates that the P5CS filaments are adjacent but not identical to the CTPS filament. Note this is the same image shown in Fig 5F.

**Figure S4.**
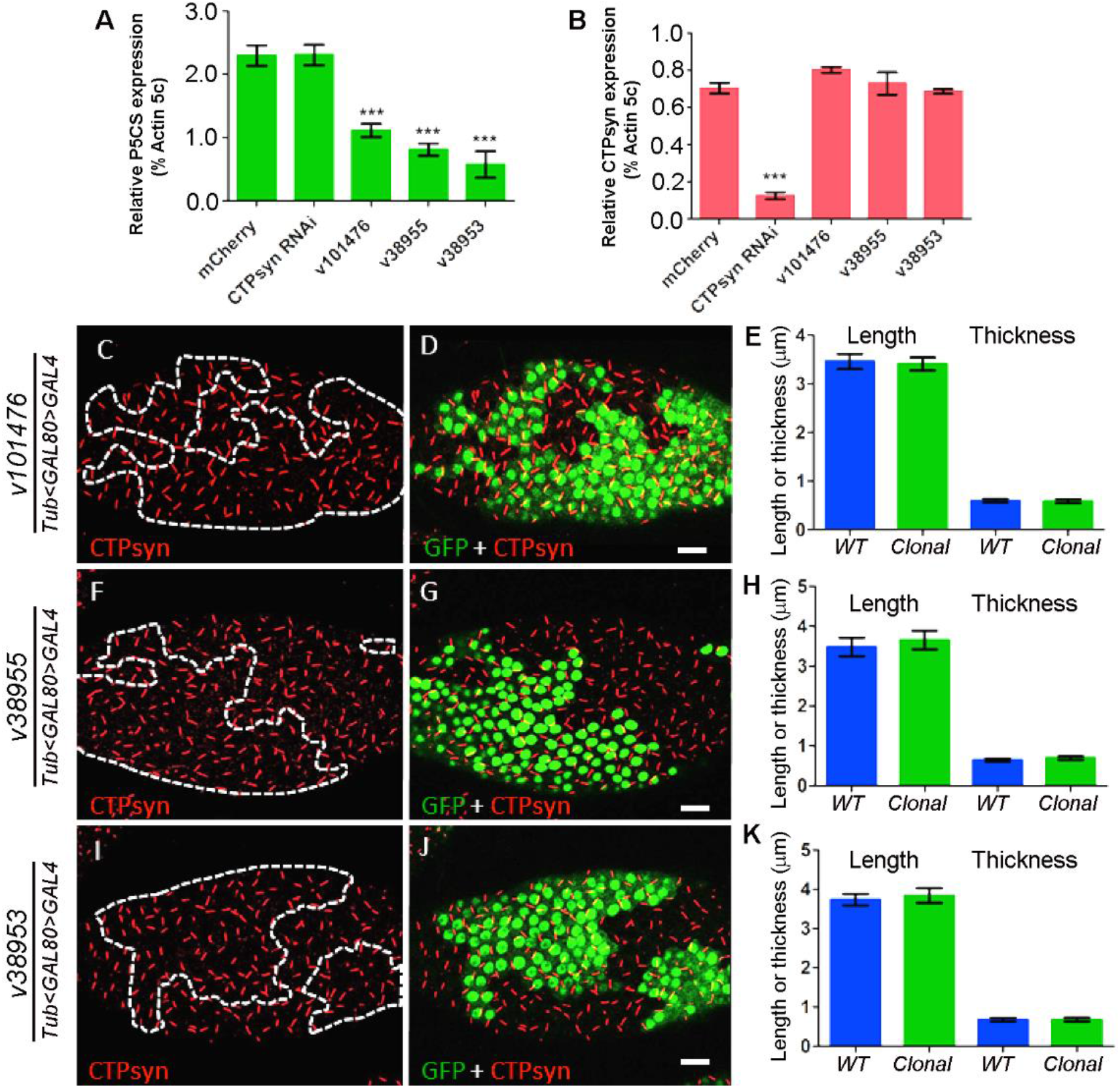
Knocking down P5CS does not affect CTPS cytoophidia. (A) P5CS transcript levels of flies after knocked down by three P5CS RNAi lines (v10176, v38955, v38953), were measured and compared with mCherry control and CTPsyn RNAi lines. Expression levels were normalized with Actin 5c levels. (B) Quantification results of CTPS mRNA levels of CTPsyn RNAi and P5CS RNAi lines in comparison with mCherry control. Error bars show SEM of three repeats. ***P<0.001. (C) and (D), (F) and (G), (I) and (J), CTPS cytoophidia images in wild-type and three P5CS RNAi induced clones (marked with GFP, outlined with dashed line). (E), (H), (K) Analysis of length and thickness of CTPS cytoophidia of wild-type and P5CS RNAi clones. Error bars show the SEM.

**Figure S5.**
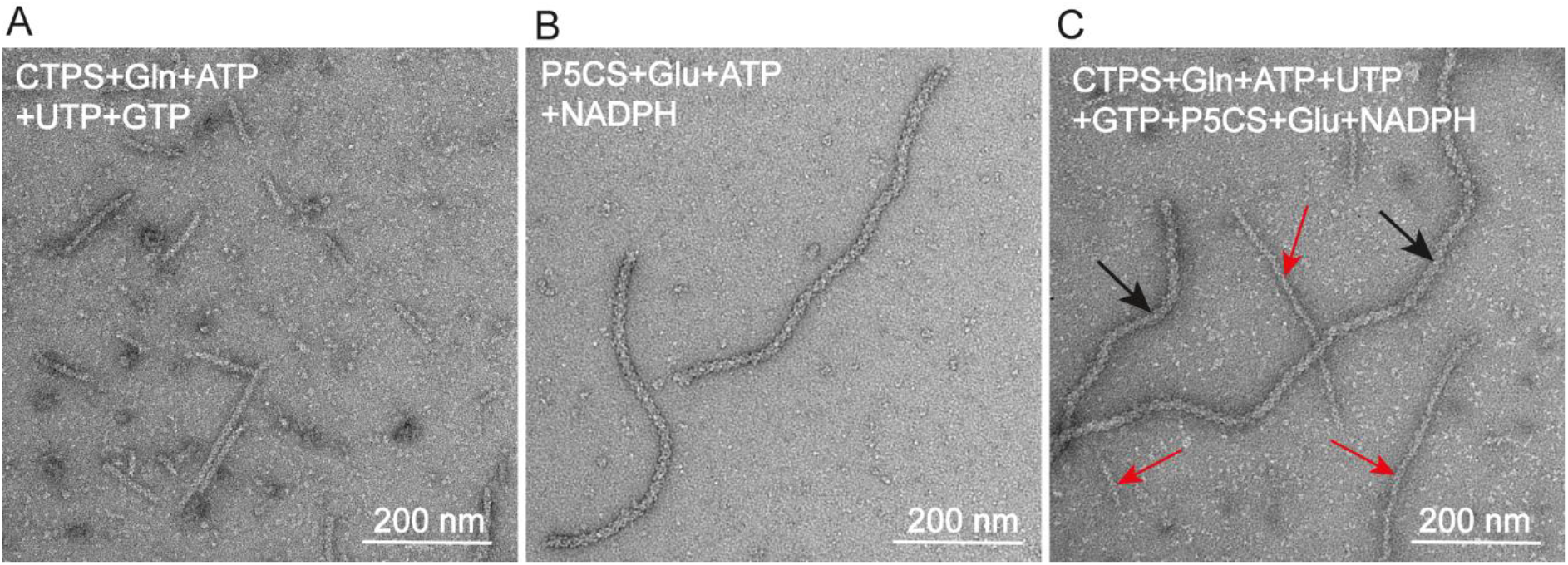
CTPS and P5CS form independent filamentous structures in vitro. (A) Negative stain images of *Drosophila* CTPS filaments in the presence of CTPS catalytical substrates. (B) *Drosophila* P5CS forms filaments in the presence of P5CS catalytical substrates. (C) Distinct filaments of CTPS and P5CS in the presence of glutamine, ATP, UTP, GTP, glutamate and NADPH. CTPS and P5CS filaments are indicated by red and black arrows, respectively.

